# The ENIGMA Toolbox: Cross-disorder integration and multiscale neural contextualization of multisite neuroimaging datasets

**DOI:** 10.1101/2020.12.21.423838

**Authors:** Sara Larivière, Casey Paquola, Bo-yong Park, Jessica Royer, Yezhou Wang, Oualid Benkarim, Reinder Vos de Wael, Sofie L. Valk, Sophia I. Thomopoulos, Matthias Kirschner, ENIGMA Consortium, Lindsay B. Lewis, Alan C. Evans, Sanjay M. Sisodiya, Carrie R. McDonald, Paul M. Thompson, Boris C. Bernhardt

**Affiliations:** Multimodal Imaging and Connectome Analysis Laboratory, McConnell Brain Imaging Centre, Montreal Neurological Institute and Hospital, McGill University, Montreal, QC, Canada; Depatment of Data Science, Inha University, Incheon, South Korea; Otto Hahn Research Group for Cognitive Neurogenetics, Max Planck Institute for Cognitive and Brain Sciences, Leipzig, Germany; INM-7, FZ Jülich, Jülich, Germany; Imaging Genetics Center, Mark & Mary Stevens Neuroimaging and Informatics Institute, Keck School of Medicine, University of Southern California, Marina del Rey, California, United States; Department of Psychiatry, Psychotherapy and Psychosomatics, Psychiatric Hospital, University of Zurich, Switzerland; http://enigma.ini.usc.edu; McConnell Brain Imaging Centre, Montreal Neurological Institute and Hospital, McGill University, Montreal, QC, Canada; McGill Centre for Integrative Neuroscience, McGill University, Montreal, QC, Canada; Department of Clinical and Experimental Epilepsy, UCL Queen Square Institute of Neurology, London, UK; Chalfont Centre for Epilepsy, Bucks, UK; Department of Psychiatry, Center for Multimodal Imaging and Genetics, University of California San Diego, La Jolla, CA, United States

**Keywords:** ENIGMA, multisite, multiscale, meta-analysis, mega-analysis, transcriptomics, connectivity

## Abstract

Among ‘big data’ initiatives, the ENIGMA (<u>E</u>nhancing <u>N</u>euroImaging <u>G</u>enetics through <u>M</u>eta-<u>A</u>nalysis) Consortium—a worldwide alliance of over 2,000 scientists diversified into over 50 Working Groups—has yielded some of the largest studies of the healthy and diseased brain. Integration of multisite datasets to assess transdiagnostic similarities and differences and to contextualize findings with respect to neural organization, however, have been limited. Here, we introduce the ENIGMA Toolbox, a Python/Matlab ecosystem for (*i*) accessing 100+ ENIGMA datasets, facilitating cross-disorder analysis, (*ii*) visualizing data on brain surfaces, and (*iii*) contextualizing findings at the microscale (*postmortem* cytoarchitecture and gene expression) and macroscale (structural and functional connectomes). Our Toolbox equips scientists with tutorials to explore molecular, histological, and network correlates of noninvasive neuroimaging markers of brain disorders. Moreover, our Toolbox bridges the gap between standardized data processing protocols and analytic workflows and facilitates cross-consortia initiatives. The Toolbox is documented and openly available at http://enigma-toolbox.readthedocs.io.

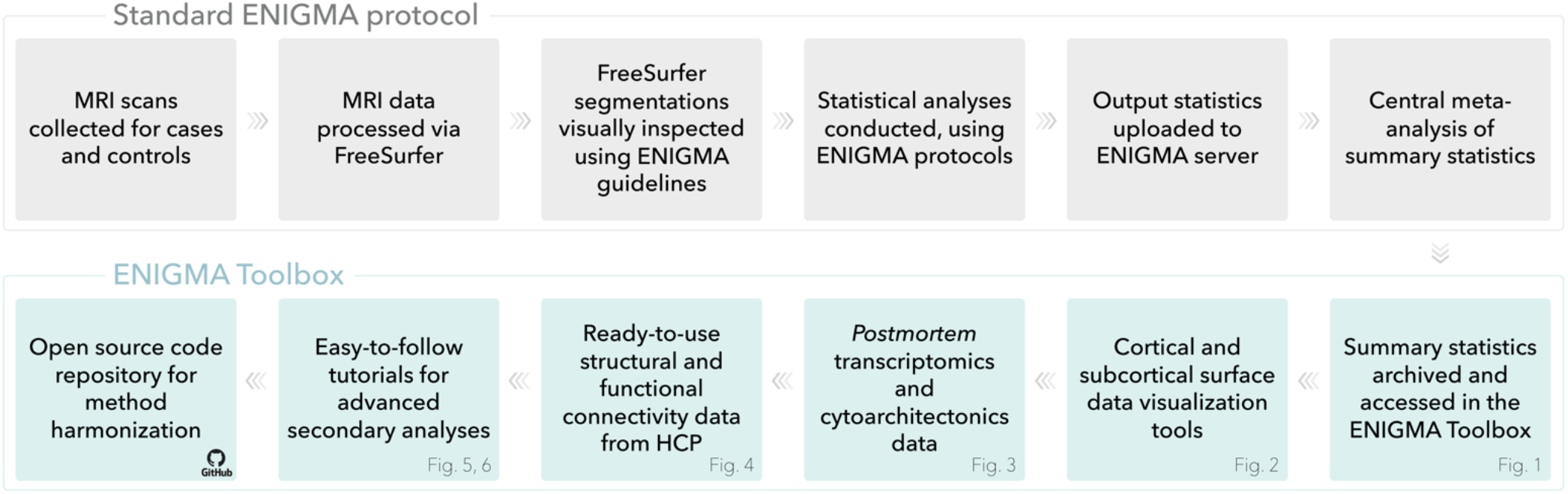

## Introduction

The ENIGMA (<u>E</u>nhancing <u>N</u>euroImaging <u>G</u>enetics through <u>M</u>eta-<u>A</u>nalysis) Consortium is one of the leading big data alliances that pools neuroimaging and genetic data from around the globe. Since its inception, ENIGMA has grown to a collaboration of thousands of scientists from over 45 countries^3^. Organized into several Working Groups (Fig. 1A), the ENIGMA Consortium has made important contributions to fundamental and clinical neuroscience, publishing some of the largest neuroimaging studies to date^4–13^. The success of the ENIGMA Consortium has heavily relied on standardized pipelines for data processing, quality control, and meta-analysis of widely used morphological measures. From these harmonized procedures, brain metrics (such as cortical thickness and subcortical volume) are extracted from raw neuroimaging data at each research center within a given Working Group and entered into site-specific linear models to test for case *vs*. control differences or to assess correlations with covariates of interest. Effect sizes and the heterogeneity of these effects across sites are then estimated via topic- and disease-specific meta-analyses, with the resulting aggregated statistical information being published and shared.

**Figure 1.**
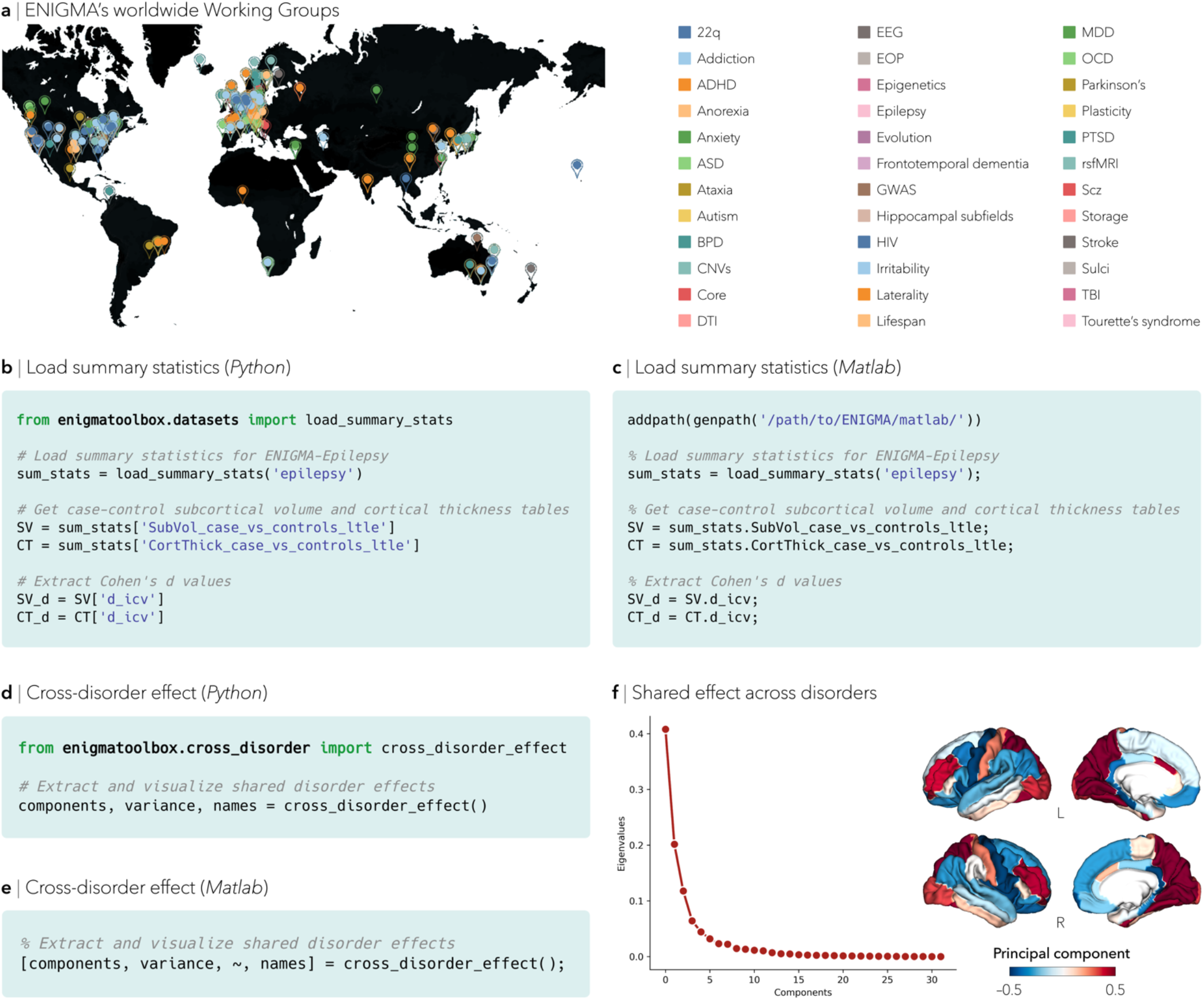
Data archiving and accessing. (**a**) World map of a subset of ENIGMA’s Working Groups (color key). Each group consists of international teams of researchers and clinicians studying major brain diseases and conditions and contributing data to the ENIGMA consortium. Case-control summary statistics from published studies are archived in the ENIGMA Toolbox and easily accessible via simple Python (**b**) and Matlab (**c**) scripts. Summary statistics from the Epilepsy Working Group is shown as an example. Minimal (**d**) Python and (**e**) Matlab code snippets to identify transdiagnostic morphometric signatures. (**f**) Eigenvalues of each component are displayed in the scree plot. The principal component underlying the shared cross-disorder effect is projected to the surface template.

Through the sharing of these site-specific brain metrics or ensuing aggregated statistical maps^14^, ENIGMA has set the stage for unprecedented analyses to compare findings across Working Groups and to contextualize findings across different scales of neural organization. Several such cross-disorder studies are underway, including a comparison of three neurodevelopmental disorders across 151 cohorts worldwide^15^, and an initiative to relate structural brain abnormalities to cell-specific profiles of gene expression^16^.

The neuroscience community, however, currently lacks standardized tools to analyze multicenter datasets beyond traditional structural MRI case-control meta- and mega-analyses. To fill this gap, we developed the ENIGMA Toolbox, an ecosystem for: (*i*) archiving, accessing, and integrating different ENIGMA-derived, or equivalently processed, datasets, (*ii*) visualizing data on cortical and subcortical surface models, and (*iii*) contextualizing neuroimaging findings across multiple scales of neural organization. We provide the ability to decode brain maps with respect to *postmortem* gene expression maps (based on microarray data from the Allen Human Brain Atlas^17^), *postmortem* cytoarchitecture (using the von Economo-Koskinas classification of cortical cytoarchitecture and derivatives from the BigBrain dataset^18^), and structural as well as functional connectome properties (from the Human Connectome Project^19^). We provide several start-to-finish tutorials to show how these workflows can offer neurobiological and system-level insights into how regional effects, for example disease-related atrophy patterns, co-vary with transcriptomic, microstructural, and macrolevel properties, similar to recently published ENIGMA studies^16, 20^.

Here, we present the ENIGMA Toolbox with ready-to-use and easy-to-follow code snippets. Our toolbox is available in Python and Matlab—two widely used languages in neuroimaging, neuroinformatics, and genetics communities—and is compatible with a range of subject-level, as well as meta- and mega-analytic datasets. Data and codes are openly accessible (http://github.com/MICA-MNI/ENIGMA) and complemented with expandable online documentation (http://enigma-toolbox.readthedocs.io).

## Results

The ENIGMA Toolbox is an ecosystem composed of three modules. Each of these modules can be stand-alone or integrated with one another, allowing for greater flexibility, adaptiveness, and continuous development. We provide thoroughly documented workflows that users can easily adapt to their own datasets.

### Data archiving and accessing

The ENIGMA Toolbox stores and accesses summary statistics from several ENIGMA Working Groups in a central repository. Given that data sharing practices can at times be challenging, in part due to privacy and regulatory protection, ENIGMA represents a practical alternative for standardized data processing and anonymized analysis of results (*i.e.*, meta-analysis) as well as the sharing of non-identifiable derivatives (*i.e.*, mega-analysis). Available datasets within our Toolbox consist of summary statistics from several ENIGMA Working Groups. The current release (*v*1.1.0) includes 100+ case-control summary statistics from eight Working Groups, including: 22.q11.2 deletion syndrome, attention deficit/hyperactivity disorder, autism spectrum disorder, bipolar disorder, epilepsy, major depressive disorder, obsessive-compulsive disorder, and schizophrenia (see METHODS). These datasets, obtained from standardized and quality-controlled protocols, represent morphological (*e.g.*, subcortical volume, cortical thickness, surface area) case-control effect sizes from previously published meta-analyses^4–6^, ^9–13^ and can be retrieved using the `load_summary_stats()` function (Fig. 1B, C) and used for secondary analyses. Summary statistics from other Working Groups will be continuously added as they are published.

As many ENIGMA groups have moved beyond meta-analysis to mega-analysis of subject-level data, the Toolbox is also compatible with subject-level data. As part of the ENIGMA Toolbox, we also provide example data from an individual site, accessible via `load_example_data()`. Alternatively, if users have generated their own summary statistics/subject-level data that adhere to ENIGMA’s harmonized processing and analysis protocols (http://enigma.ini.usc.edu/protocols/), or have in-house parcellated or vertexwise datasets, they can import their own data locally and take advantage of every function our Toolbox has to offer. Online tutorials, including those related to data importation/exportation, are suited for all dataset types.

To yield novel insights into brain structural abnormalities that are common or different across disorders, available summary statistics can also be harnessed to conduct cross-disorder analyses. From the `cross_disorder_effect()` function, users can explore shared and disease-specific morphometric signatures with two different approaches: (*i*) by applying a principal component analysis (PCA) to any combination of disease-specific summary statistics, resulting in shared latent components that can be used for further analysis (Fig. 1D–F), and (*ii*) by systematically cross-correlating patterns of brain structural abnormalities with every other set of available summary statistics, resulting in a correlation matrix (Fig. S1A–C). Users can also upload local summary statistics, or equivalently processed data, for inclusion in the PCA or comparison against our database.

### Cortical and subcortical surface data visualization

The ENIGMA Toolbox provides functions to visualize cortical and subcortical data on surface models and generate publication-ready figures (Fig. 2A, B). To illustrate the `plot_cortical()` and `plot_subcortical()` functions, we projected cortical and subcortical gray matter atrophy in individuals with left focal epilepsy relative to healthy controls to the surface templates (Fig. 2C). Beyond the mapping of gray matter atrophy, our surface visualization function is compatible with any neuroimaging data type, parcellation, and surface templates. Mapping to and from brain parcellations and the vertex-wise surface space can also be easily achieved using the `parcel_to_surface()` and `surface_to_parcel()` functions.

**FIGURE 2.**
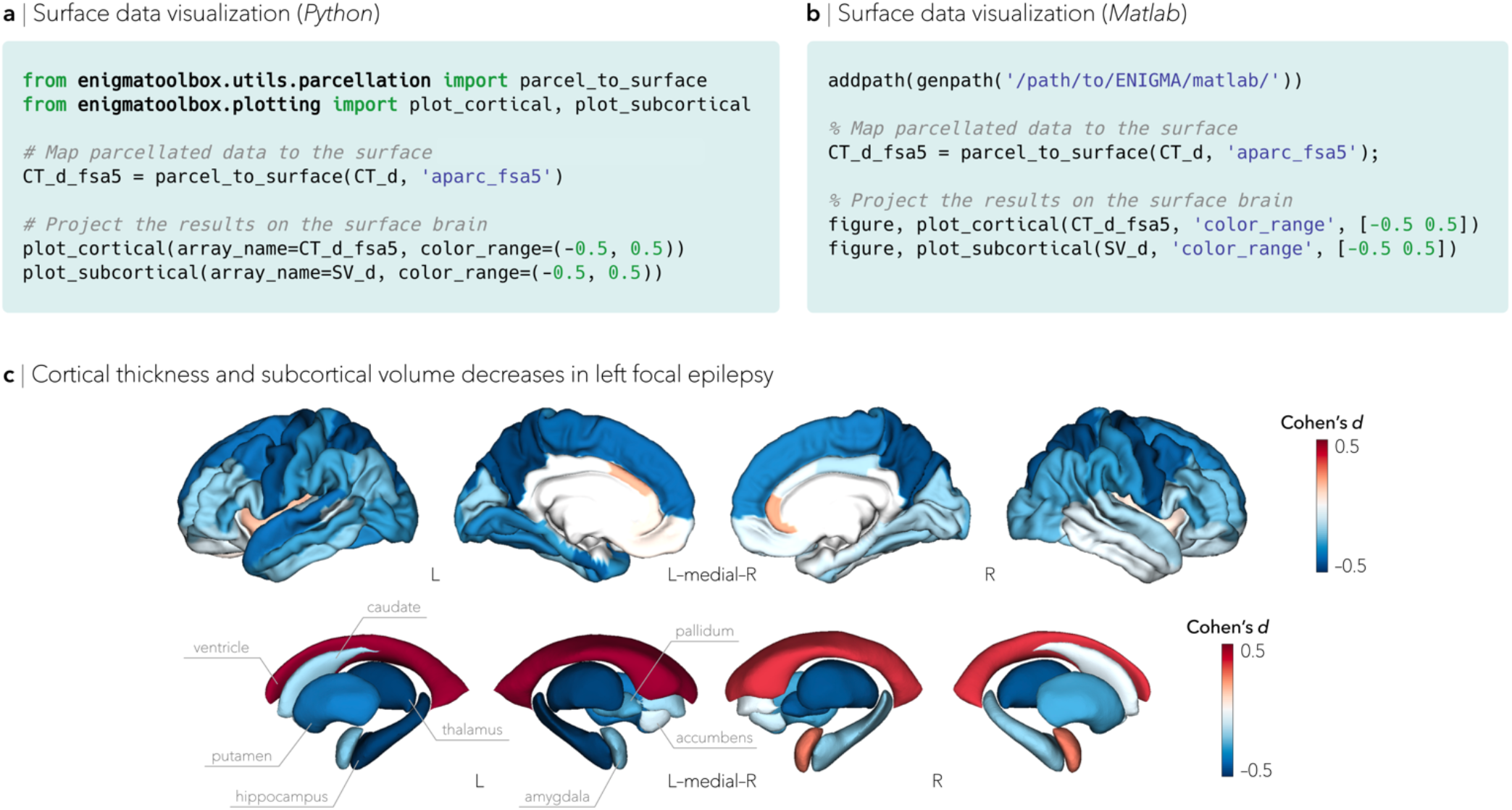
Cortical and subcortical surface visualization. Minimal (**a**) Python and (**b**) Matlab code snippets for plotting cortical thickness and subcortical volume deficits in individuals with left focal epilepsy. (**c**) Summary statistics (Cohen’s *d*) comparing individuals with left focal epilepsy to healthy controls are projected to the surface templates. Profound gray matter atrophy can be visually appreciated in bilateral precentral gyrus, precuneus, thalamus, as well as left mesiotemporal regions, including the hippocampus.

### Multiscale contextualization

#### Transcriptomics data

The emergence of open databases for human transcriptomics yields new opportunities to associate macroscale neuroimaging findings with spatial variations at the molecular scale^17, 21–23^. The Allen Institute for Brain Science released the Allen Human Brain Atlas (AHBA)—a brain-wide gene expression atlas comprising microarray-derived measures from over 20,000 genes sampled across 3,702 spatially distinct tissue samples^17^. Using the abagen toolbox^24^ and following guidelines established by Arnatkevic̆iūtė et al.^21^, this large expression dataset was collapsed into cortical and subcortical regions of interest and combined across donors^1, 21, 24^. Genes that were consistently expressed across donors (*r*≥0.2, *n*_genes_=12,668), can be easily obtained using the `fetch_ahba()` function (Fig. 3A–C). Additional details on processing and analytical choices, including parcellation compatibility and stability thresholds, are provided in the Methods.

**FIGURE 3.**
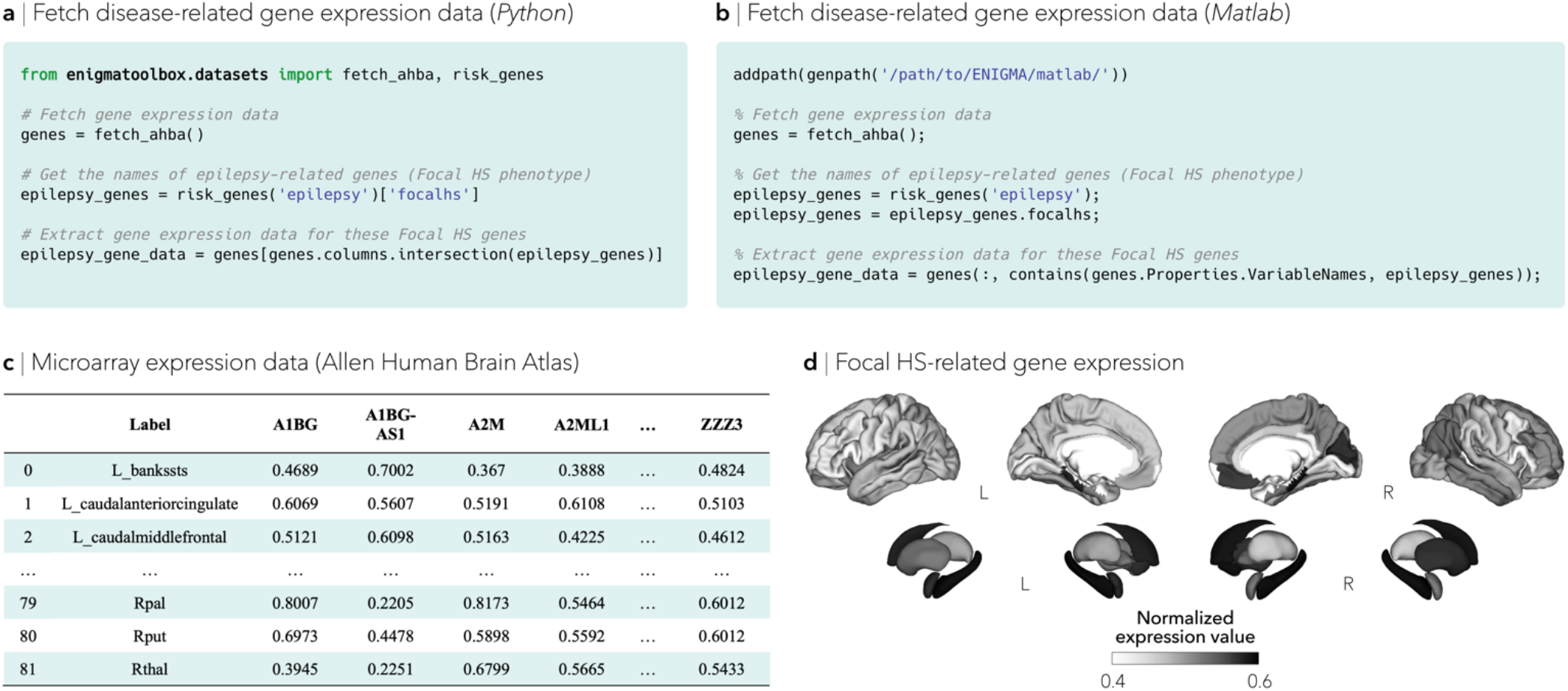
ENIGMA-friendly gene co-expression data. Minimal (**a**) Python and (**b**) Matlab code snippets to fetch disease-related gene co-expression data. Genes related to epilepsy, more specifically focal hippocampal sclerosis (Focal HS), are extracted as an example. (**c**) The complete microarray expression data can be easily accessed from the ENIGMA Toolbox and contains data from over 15,000 genes. (**d**) Disease-related gene co-expression data can be mapped to the surface templates; here, we displayed the average expression levels of Focal HS genes on the cortical and subcortical surface templates.

#### BigBrain data

The BigBrain, an ultra-high-resolution 3D reconstruction of a sliced and stained human brain^18^, has supported quantitative analysis of cytoarchitecture. As part of the ENIGMA Toolbox, we characterized regional cytoarchitecture using statistical moments of staining profiles (see Methods)^25^. Specifically, studying the mean intracortical staining across the mantle allows inferences on overall cellular density, whereas analysis of profile skewness indexes the distribution of cells across upper and lower layers of the cortex—a critical dimension of laminar differentiation.

In addition to statistical moments, prior work has demonstrated robust evidence for a principal gradient of gradual cytoarchitectural variation running from primary sensory to limbic areas, mirroring spatial trends in laminar differentiation and cytoarchitectural variations^26–28^. To expand our BigBrain contextualization module, we also incorporated this microstructural similarity gradient to describe a sensory-fugal transition in intracortical microstructure (see Methods). Stratifying cortical findings relative to this gradient could, for example, test whether patterns of changes are conspicuous in cortices with marked laminar differentiation (*e.g.*, sensory and motor cortices) or in those with subtle laminar differentiation (*e.g.*, limbic cortices).

#### Cytoarchitectural types

To further describe microscale cortical organization, the ENIGMA Toolbox includes a digitized parcellation of the von Economo and Koskinas cytoarchitectonic map of the human cerebral cortex^2, 29^. From this mapping, five different structural types of cerebral cortex are recorded: *i*) agranular (thick with large cells but sparse layers II and IV), *ii*) frontal (thick but not rich in cellular substance, visible layers II and IV), *iii*) parietal (thick and rich in cells with dense layers II and IV but small and slender pyramidal cells), *iv*) polar (thin but rich in cells, particularly in granular layers), and *v*) granular or koniocortex (thin but rich in smalls cells, even in layer IV, and a rarified layer V)^30^.

#### Connectivity data

Neuroimaging, particularly with functional and diffusion MRI, has become a leading tool to characterize human brain network organization *in vivo* and to identify network alterations in brain disorders. Although ongoing efforts in ENIGMA and beyond are beginning to coordinate analyses of resting-state functional^31–33^ and diffusion^34–36^ MRI data, connectivity measures remain sparse within the consortium. As an alternative, our Toolbox leverages high-resolution structural (derived from diffusion-weighted tractography) and functional (derived from resting-state functional MRI) connectivity data from a cohort of unrelated healthy adults from the Human Connectome Project (HCP)^19^. Preprocessed cortico-cortical, subcortico-cortical, and subcortico-subcortical functional and structural connectivity matrices can be easily retrieved using the `load_fc()` and `load_sc()` functionalities (Fig. 4A–C). Connectivity matrices were parcellated according to several atlases, including the Desikan-Killiany atlas, and can thus be readily combined with any ENIGMA-derived, or other parcellated, datasets1. Details on subject inclusion, data preprocessing, and matrix generation are provided in the Methods.

**FIGURE 4.**
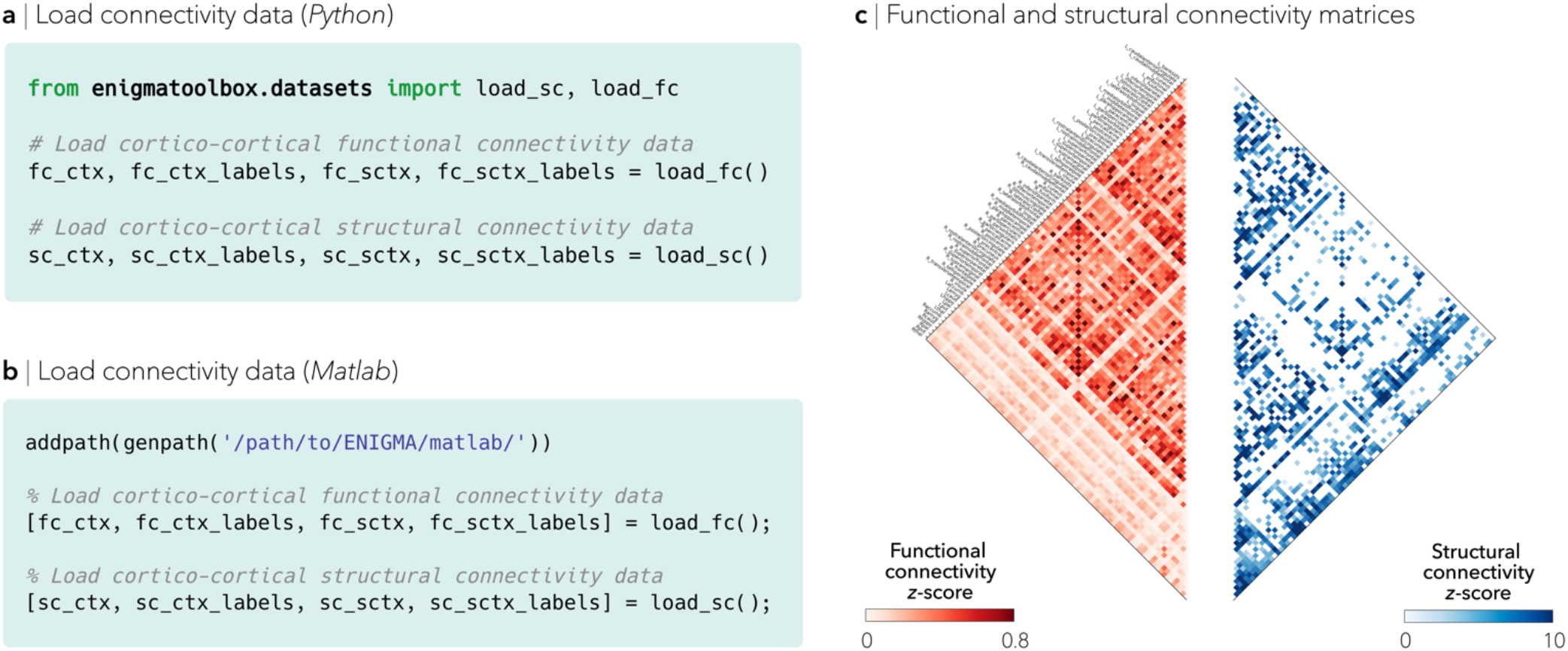
High-resolution connectivity data sharing and exploiting. Minimal (**a**) Python and (**b**) Matlab code snippets to load preprocessed functional and structural connectivity matrices. (**c**) Available connectivity data within the ENIGMA Toolbox include cortico-cortical, subcortico-cortical, and subcortico-subcortical connectivity matrices. Matrices are unthresholded and parcellated according to the Desikan-Killiany atlas^1^.

### Analytical workflows

As of the current release (*v*1.1.0), the ENIGMA Toolbox comprises two neural scales for the contextualization of findings: (*i*) using microscale properties, namely gene expression and cytoarchitecture, and (*ii*) using macroscale network models, such as regional hub susceptibility analysis and disease epicenter mapping. Albeit often restricted to individual site studies, similar approaches have been used to study microstructural organization in healthy^26, 37, 38^ and diseased^39^ brains, and to model network-level patterns of disease-related atrophy^40–42^. To ease programming and maximize transparency, analytic workflows are accompanied by comprehensive tutorials and visual assessment checkpoints. As proofs of concept, we demonstrate ready-to-use, easy-to-follow, and validated secondary analysis tutorials to relate patterns of gray matter atrophy in individuals with left focal epilepsy to transcriptomic, histological, and normative connectome properties.

#### Transcriptomics contextualization of findings

Motivated by the growing body of research relating gene expression to diverse properties of macroscale brain organization, Toolbox users can import the AHBA microarray expression dataset and visualize brain maps of gene expression levels. This tool can also be used to identify genes that are spatially correlated with a given brain map (*e.g.*, a disease-related atrophy map). Moreover, based on reports of recently published genome-wide association studies (GWAS)^43–49^, users can extract the most likely genes associated to significant genome-wide loci across a range of disorders. From these sets of genes, users can generate disease-specific gene expression maps to contextualize and decode neuroimaging findings with transcriptomics data. To illustrate the `risk_genes()` function, we selected genes related to focal epilepsy with hippocampal sclerosis^47^ and displayed their average expression levels on the cortical and subcortical surface templates (Fig. 3D). In this example, we can observe higher gene co-expression levels of epilepsy-related genes in mesiotemporal lobe regions, overlapping with regions of profound atrophy in individuals with left focal epilepsy. Related efforts are underway to map the effects of common (single nucleotide) variants in the genome on brain structure, using visualization platforms such as ENIGMA-Vis (https://enigma-brain.org/enigmavis/)^50^ and the Oxford Brain Imaging Genetics browser (http://big.stats.ox.ac.uk/).

#### Histological contextualization of findings

Applied conjointly with ENIGMA datasets, BigBrain-derived intracortical profile information (*i.e.*, the statistical moments and the principal gradient of microstructure differentiation) offers two complementary approaches to situate cortical findings with respect to histological findings. For the former approach, users can feed a thresholded cortical map—for instance areas of significant structural abnormalities—into the `bb_moments_raincloud()` function to contextualize macroscale features with respect to microstructural profiles (*e.g.*, cellular density, cellular distribution asymmetry). This quantitative approach has been used to guide boundary definition and fingerprint cytoarchitecture in studies of *postmortem* data^51, 52^. Alternatively, thresholded (or unthresholded) cortical maps can be fed into the `bb_gradient_plot()` function, which discretizes the principal microstructure similarity gradient into five equally-sized bins and averages surface findings within each gradient bin. From this, neuroimaging data can be embedded into the gradient space, allowing users to make inferences about the underlying microstructural hierarchy of, for instance, atrophied regions. Contextualizing gray matter atrophy patterns in left focal epilepsy with respect to histological properties, we were able to demonstrate that atrophy predominantly affected cortical regions with greater, and more evenly distributed, cellular densities across upper and lower layers of the cortex (Fig. S2A–D)—areas located towards the sensory apex of the cytoarchitectural gradient (Fig. S3A–D).

#### Cytoarchitectonic contextualization of findings

Integration of cytoarchitecture with *in vivo* neuroimaging can consolidate microstructural differentiation with macroscale-level effects. As part of the Toolbox, users can also leverage a digitized cytoarchitectonic atlas from seminal *postmortem* work by von Economo and Koskinas^2^. From the `economo_koskinas_spider()` function, users can apply this well-established decomposition to summarize cortex-wide effects (*e.g.*, a disease-related atrophy map) and assess relationships to distinct cytoarchitectonic classes. To ease interpretability, cytoarchitectonic classification of findings are also displayed in a spider plot. To highlight how atrophy can vary with distinct architectonic cortical types, case-control Cohen’s *d* measures, representing atrophy in left focal epilepsy, were averaged within each von Economo and Koskinas cytoarchitectonic class. Projecting the results in a radar (spider) plot revealed that the agranular cortex was most affected by atrophy in left focal epilepsy (Fig. S4A–C).

#### Hub susceptibility model

Normative structural and functional connectomes hold valuable information for relating macroscopic brain network organization to patterns of disease-related atrophy (Fig. 5A, B). Prior work studying network underpinnings of morphological abnormalities in neurodegenerative and psychiatric disorders has demonstrated that hubs (*i.e.*, brain regions with many connections) typically show greater atrophy than locally-connected peripheral nodes^53, 54^. Within the ENIGMA Epilepsy Working Group, we recently tested this hypothesis using data from 1,021 individuals with epilepsy and 1,564 healthy controls, and also showed that atrophy preferentially colocalized with highly interconnected hub regions in the common epilepsies^20^. A series of follow-up ENIGMA studies are currently underway to assess these network-level effects in other disorders.

**FIGURE 5.**
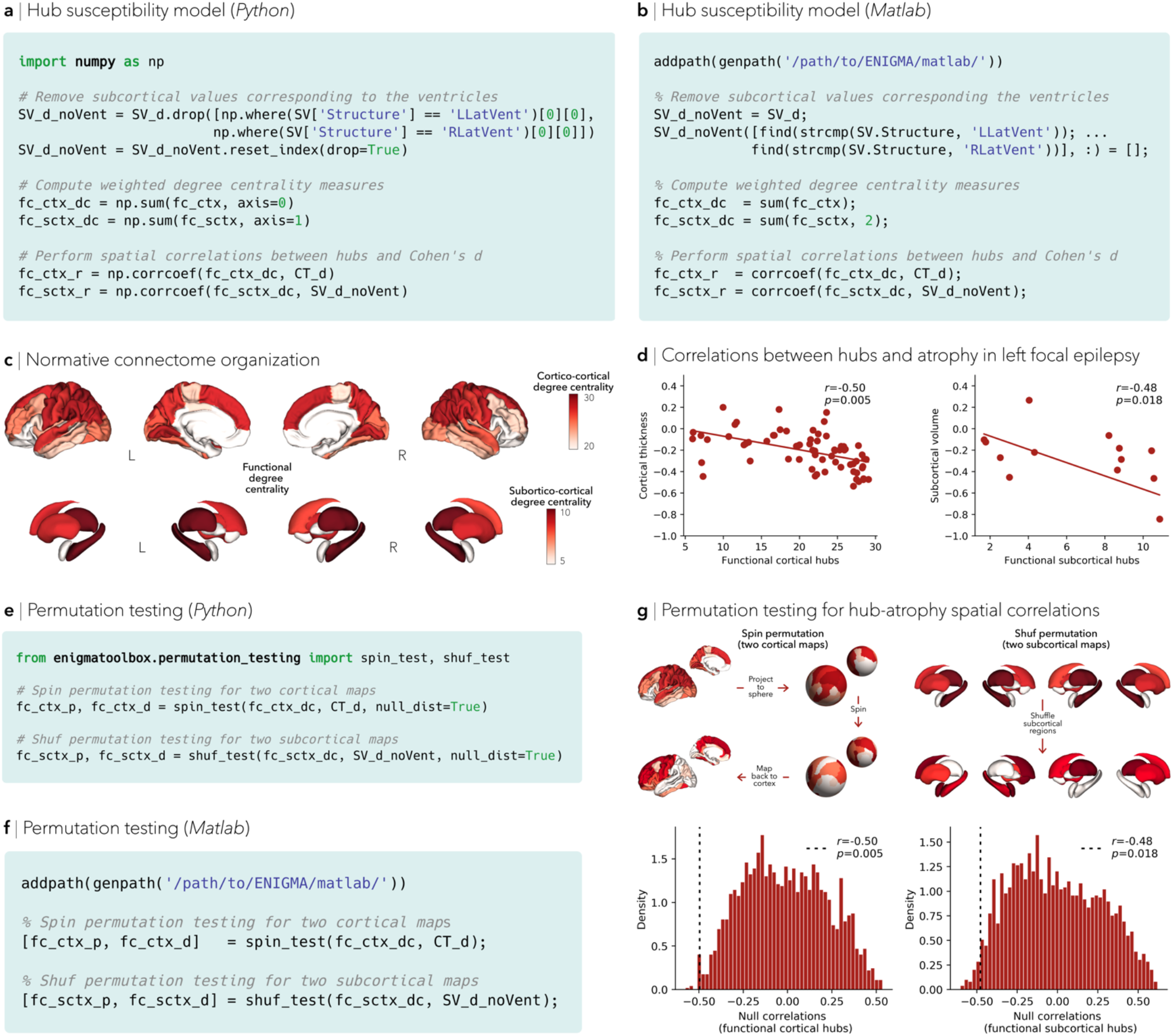
Advanced analytical workflows: hub susceptibility models and spin permutation testing. Minimal (**a**) Python and (**b**) Matlab code snippets to assess relationships between hubs and patterns of atrophy. (**c**) Functional degree centrality, derived from the HCP dataset, was used to identify the spatial distribution of hub regions. (**d**) Strong negative associations between cortical thickness/subcortical volume Cohen’s *d* values and cortico-/subcortico-cortical degree centrality were observed, indicating that atrophy preferentially colocalized with hub regions. Minimal (**e**) Python and (**f**) Matlab code snippets to assess statistical significance of two surface maps while preserving spatial autocorrelation. (**g**) A schematic of the spin and shuf permutation frameworks for cortical and subcortical maps, respectively. The null distributions of correlations are shown; the empirical (*i.e.*, original) correlation coefficients and associated spin permuted p-values are indexed by the dashed line.

Building on the above-described functions, we can first derive weighted degree centrality maps from functional (or structural) connectivity data by computing the sum of all weighted cortico- and subcortico-cortical connections for every region, with higher degree centrality denoting hub regions (Fig. 5C). Spatial similarity between atrophy patterns (obtained from individuals with left focal epilepsy as an example) and hub distributions can then be compared through correlation analysis (and statistically assessed via spin permutation tests; see below), revealing that profound atrophy implicates functional cortico- and subcortico-cortical hubs more strongly than nonhub regions (Fig. 5D).

#### Disease epicenter model

To further investigate whether disease-related morphological abnormalities (*e.g.*, atrophy) follow overarching principles of connectome organization, one can identify disease epicenters. Disease epicenters are regions whose functional and/or structural connectivity profile spatially resembles a given disease-related atrophy map^20, 40–42^. Hence, disease epicenters can be identified by spatially correlating every region’s healthy functional and/or structural connectivity profiles to whole-brain atrophy patterns in a given disease. This approach must be repeated systematically across the whole brain, assessing the statistical significance of the spatial similarity of every region’s functional and/or structural connectivity profiles to disease-specific abnormality maps with spatial permutation tests. Cortical and subcortical epicenter regions can then be identified if their connectivity profiles are significantly correlated with the disease-specific abnormality map. Regardless of its atrophy level, a cortical or subcortical region could potentially be an epicenter if it is (*i*) strongly connected to other high-atrophy regions and (*ii*) weakly connected to low-atrophy regions. Moreover, disease epicenters do not necessarily represent hub regions, but may rather be connected to them (*i.e.*, so-called ‘feeder nodes’, which directly link peripheral nodes to hubs). As in prior work^20^, this approach suggests that patterns of atrophy in left focal epilepsy are anchored to the connectivity profiles of mesiotemporal regions (Fig. S5A–C).

#### Statistical comparison of spatial maps

The intrinsic spatial smoothness of brain maps—where data from neighboring regions are not statistically independent of each other—violates the underlying assumptions of several inferential statistical tests and consequently inflates the apparent significance of their spatial correlation, unless more sophisticated tests are used. To overcome these shortcomings and minimize Type I error, our Toolbox includes non-parametric spatial permutation models to assess statistical significance while preserving the spatial autocorrelation of brain maps^55, 56^. With such functionality, for instance `spin_test()`, the spatial coordinates of the surface data are projected onto the surface spheres and randomly rotated to generate surface maps with randomized topography but identical spatial autocorrelation. The empirical (*i.e.*, real) correlation coefficient can then be compared against the null distribution determined by the ensemble of correlation coefficients comparing spatially permuted surface maps. Spatial correspondence between two subcortical surface maps can be examined via a standard non-parametric null model, namely `shuf_test()`, where subcortical labels are randomly shuffled as opposed to being projected onto spheres. To illustrate these functions, we assessed the statistical significance of cortical and subcortical hub-atrophy correlations in left focal epilepsy (Fig. 5E, F) and displayed the empirical correlation coefficients onto the null distributions of permuted correlations (Fig. 5G).

## Discussion

The ENIGMA Toolbox is an integrated ecosystem that dovetails with ENIGMA’s standardized data processing and meta-analysis strategies for integration, visualization, and contextualization of multisite results. Our Toolbox relies on a simple but efficient codebase for exploring and analyzing big data, aiming to facilitate and homogenize follow-up analyses of ENIGMA, or other, MRI datasets around the globe. At its core, the Toolbox is composed of three modules for (*i*) archiving, accessing, and integrating case-control meta-analytic datasets from specialized Working Groups, (*ii*) cortical and subcortical surface data visualization, and (*iii*) contextualizing findings based on transcriptomics, cytoarchitecture, and connectivity data. Owing to its comprehensive tutorials, detailed functionality descriptions, and visual reports, our Toolbox is accessible to researchers and clinicians without extensive programming expertise within and beyond ENIGMA itself.

Enriching *in vivo* morphological correlates with *postmortem* microstructural information can deepen our understanding of the molecular and cellular underpinnings of healthy and diseased brain organization. To illustrate such microscale contextualization, we provide several tutorial examples that reference ENIGMA-type maps of gray matter atrophy in individuals with epilepsy against *postmortem* gene co-expression and histological measures. Based on the gene expression atlas from the Allen Institute for Brain Science17, which compiles information on transcription of thousands of genes across the adult brain, users can compare spatial patterns of brain-wide gene expression to MRI-derived neuroanatomy measures. Importantly, microarray expression data within our Toolbox were processed according to recommendations for best practice summarized in Arnatkevic̆iūtė and colleagues using the open-access abagen toolbox (https://abagen.readthedocs.io/en/stable/)^24^, facilitating comparisons of findings across studies. Prior neuroimaging studies have already identified specific transcriptomic signatures of cortical morphometry in early brain development^57^, cortical anatomy changes in youth with known genomic dosage variations^58^, myeloarchitectural development in adolescence^59^, and autism pathophysiology^60^, bridging microstructural and macroscopic scales of brain organization (for a review, see ^22^). Similarly, in a large-scale collaborative effort involving six ENIGMA Working Groups, Patel and colleagues correlated brain-wide cell-specific gene expression with group differences in cortical thickness, revealing shared neurobiological processes that underlie morphological phenotypes of multiple psychiatric disorders^16^. Moreover, as part of our toolbox, we also made the digital cell body-stained BigBrain^18^ and von Economo and Koskinas cytoarchitectural atlas^2, 29^ easily accessible and compatible with various neuroimaging parcellations. From the 3D histological BigBrain, several groups have made promising headway into mapping the patterns of cortical laminar architecture^61^, exploring histological underpinnings of MRI-based thickness gradients in sensory and motor cortices^62^, and identifying a sensory-fugal axis of microstructural differentiation^26^. On the other hand, histological atlases, such as the one by von Economo and Koskinas, are invaluable for linking brain microstructure to functional localization^30^. The digitized parcellation of von Economo and Koskinas cytoarchitectural types, thus, enables users to speculate on the underlying cytoarchitectural composition of, for instance, structurally abnormal areas in specific diseases. When combined, transcriptomic and cytoarchitectonic decoding can embed neuroimaging findings in a rich neurobiological context and yield potentially novel insights into the etiology of several brain disorders.

At the macroscopic level, network connectivity offers a vantage point to quantify brain reorganization in diseases that are increasingly being conceptualized as network disorders^63–65^. To exemplify the potential of macroscale network modeling, our Toolbox provides detailed tutorials on how to relate neuroimaging surface maps to normative connectome properties derived from functional and diffusion MRI. Building on prior neurodegenerative^42, 53^ and psychiatric^66^ research, as well as recent work from our group^20^, Toolbox users can build hub susceptibility models to assess the vulnerability of highly connected network hubs to disease-related effects. Given their role in integrative processing, as well as their high topological value and biological cost, hub regions have been hypothesized to be preferentially susceptible to diverse pathological perturbations^67, 68^. Consequently, hub decoding represents an invaluable tool to interrogate whether heightened connectivity, atrophy, or metabolism properties in hubs relate to disease-specific processes. Complementing the hub susceptibility approach, another network modeling functionality available within our Toolbox is disease epicenter mapping, an analysis technique originally developed to study the spread of atrophy in neurodegenerative diseases^69^. Subsequent *in vivo* neuroimaging research provided evidence that network connectivity may herald patterns of gray matter atrophy in schizophrenia^40^, Parkinson’s disease^42, 70^, and frontotemporal dementia syndromes^41^, each linking greater atrophy to the connectivity profiles of distinct epicenters. Also applied to the common epilepsies as part of an ENIGMA-Epilepsy secondary project^20^, this approach identified mesiotemporal epicenters in temporal lobe epilepsy and subcortico-cortical epicenters in generalized epilepsy, regions known to be involved in pathophysiology of each syndrome^71–75^. Combined, these two network models can significantly advance our understanding of how connectome architecture relates to morphological abnormalities across a range of disorders. Indeed, as with the microscale contextualization, our macroscale hub- and epicenter-based analytic pipelines can be applied to any neuroimaging datasets. Network models can be further enriched with microstructural properties to inject multiscale information into cortical and subcortical morphometric findings^76^.

To enhance interpretability and avoid ‘black-box’ solutions, datasets, codes, and functionalities within our Toolbox are openly accessible and thoroughly documented. We hope that these efforts accelerate research and increase reliability and reproducibility. Notably, the modular architecture of our Toolbox allows for continuous development of analytical functionalities and tutorials. Future planned releases are poised to embrace new scientific approaches as they are published, adapting to new datasets (*e.g.*, PsychEncode consortium^77^), modalities (*e.g.*, resting-state functional MRI), and analytic pipelines (*e.g.*, structural covariance network analysis). Extension of the ENIGMA Toolbox to increase versatility of secondary analyses to additional brain parcellations, as well as vertex- and voxel-wise space, is already part of the development roadmap and will be updated to accommodate users’ requests. Integration of analytic methods from users around the globe is supported and encouraged to maximize the contribution, reusability, and adaptability of any neuroimaging datasets.

In closing, by bridging the gap between pre-established data processing protocols and several analytic workflows, we hope that the ENIGMA Toolbox facilitates neuroscientific contextualization of results and cross-consortia initiatives. We are eager for researchers and clinicians to test hypotheses beyond traditional case-control comparisons. We hope that our platform will lead to novel and harmonized analyses in global neuroimaging initiatives.

## Materials and methods

### Code data availability

All code used for data analysis and visualization is available on GitHub (http://github.com/MICA-MNI/ENIGMA). The ENIGMA Toolbox Python package relies on the following open-source dependencies: Matplotlib^78^, NiBabel^79^, nilearn^80^, Numpy^81, 82^, pandas^83^, seaborn^84^, Scikit-learn^85^, SciPy^86^, and VTK^87^. Users seeking help are encouraged to subscribe and post their questions to the ENIGMA Toolbox mailing list at https://groups.google.com/g/enigma-toolbox.

### ENIGMA data description

#### Meta-analytical comparisons (summary statistics)

ENIGMA’s standardized protocols for data processing, quality assurance, and meta-analysis of individual subject data were conducted at each site (http://enigma.ini.usc.edu/protocols/imaging-protocols/). For site-level meta-analysis, all research centres within a given specialized Working Group tested for case *vs*. control differences using linear models, where diagnosis (*e.g.*, healthy controls *vs*. individuals with epilepsy) was the predictor of interest, and subcortical volume, cortical thickness, or surface area of a given brain region was the outcome measure. Case-control differences were computed across all regions using either Cohen’s *d* effect sizes or *t*-values, after adjusting for different combinations of age, sex, dataset/scan site/scanner differences, intracranial volume (IVC), and intelligence quotient (IQ) effects (see Table S1 and online documentation for disease-specific models). Across-site random-effects meta-analyses of Cohen’s *d*/*t*-values were then performed for each of the cortical and subcortical region. These ENIGMA summary statistics can be retrieved from the ENIGMA Toolbox and contains the following data: effect sizes for case-control differences (d_icv), standard error (se_icv), lower bound of the confidence interval (low_ci_icv), upper bound of the confidence interval (up_ci_icv), number of controls (n_controls), number of patients (n_patients), observed p-values (pobs), false discovery rate (FDR)-corrected p-value (fdr_p).

#### Individual site or mega-analytic data

Functionalities and tutorials within the ENIGMA Toolbox are generalizable to individual site or mega-analysis datasets, which pool individual-level data. Due to restrictions of individual-level data transfer, however, we provide an ENIGMA-derived example dataset that includes fully anonymized data from ten healthy controls (7 females, age±SD=33.3±8.8 years) and ten individuals with drug-resistant temporal lobe epilepsy (7 females, age±SD=39.8±14.8 years). The Ethics Committee of the Montreal Neurological Institute and Hospital approved the study. Written informed consent, including a statement for open sharing of collected data, was obtained from all participants. As per ENIGMA-Epilepsy protocols, users can fetch covariate information (subject ID, diagnosis, sub-diagnosis, handedness, age at onset, duration of illness, and ICV), regional volumetric data (from 12 subcortical regions—namely bilateral accumbens, amygdala, caudate, pallidum, putamen, and thalamus, bilateral hippocampus, and bilateral ventricles), and cortical thickness and surface area from every Desikan-Killiany cortical region^1^. Given the standardized ENIGMA format, users can easily replace our example dataset with any other individual site or mega-analysis datasets of their own.

#### Compatibility with other datasets

To increase generalizability and usability, every function within the ENIGMA Toolbox is compatible with any neuroimaging data parcellated according to the Desikan-Killiany^1^, Glasser^88^, and Schaefer^89^ parcellations (other parcellations will be added upon request).

### Transcriptomics data and contextualization

As part of the ENIGMA Toolbox, users can fetch and manipulate preprocessed microarray expression data collected from six human donor brains and released by the Allen Institute for Brain Sciences^17^. Microarray expression data were first generated using abagen^24^, a toolbox that provides reproducible workflows for processing and preparing gene co-expression data according to previously established recommendations^21^; preprocessing steps included intensity-based filtering of microarray probes, selection of a representative probe for each gene across both hemispheres, matching of microarray samples to brain parcels from the Desikan-Killiany^1^, Glasser^88^, and Schaefer^89^ parcellations, normalization, and aggregation within parcels and across donors. Moreover, genes whose similarity across donors fell below a threshold (*r*<0.2) were removed, leaving, for instance, a total of 12,668 genes for analysis using the Desikan-Killiany atlas. To accommodate users, we also provide unthresholded gene datasets with varying stability thresholds (*r*≥0.2, *r*≥0.4, *r*≥0.6, *r*≥0.8) for every parcellation (https://github.com/saratheriver/enigma-extra).

ENIGMA Toolbox users can furthermore query pre-defined lists of disease-related genes (obtained from several recently published GWAS), including gene sets for attention deficit/hyperactivity disorder (*n*_genes_=26)^43^, autism spectrum disorder (*n*_genes_=30)^44^, bipolar disorder (*n*_genes_=30)^45^, depression (*n*_genes_=269)^46^, common epilepsies (*n*_genes_=21)^47^, schizophrenia (*n*_genes_=213)^48^, and Tourette’s syndrome (*n*_genes_=58)^49^. These gene sets can be subsequently mapped to cortical and subcortical regions using the Allen Human Brain Atlas^17^ and projected to surface templates using our surface visualization tools.

### BigBrain data and contextualization

BigBrain is a ultra-high resolution, 3D volumetric reconstruction of a *postmortem* Merker-stained and sliced human brain from a 65-year-old male, with specialized pial and white matter surface reconstructions (obtained via the open-access BigBrain repository: https://bigbrain.loris.ca/main.php)^18^. The *postmortem* brain was paraffin-embedded, coronally sliced into 7400 20μm sections, silver-stained for cell bodies^90^, and digitized. A 3D reconstruction was implemented with a successive coarse-to-fine hierarchical procedure^91^, resulting in a full brain volume. For the ENIGMA Toolbox, we used the highest resolution full brain volume (100μm isotropic voxels), then generated 50 equivolumetric surfaces between the pial and white matter surfaces. The equivolumetric model compensates for cortical folding by varying the Euclidean distance between pairs of intracortical surfaces throughout the cortex, thus preserving the fractional volume between surfaces^92^. Next, staining intensity profiles, representing neuronal density and soma size by cortical depth, were sampled along 327,684 surface points in the direction of cortical columns.

Following seminal histological work^51, 52, 59^, we characterized vertex-wise cytoarchitecture by taking two central moments of the staining intensity profiles (mean and skewness). Finally, the Desikan-Killiany atlas was nonlinearly transformed to the BigBrain histological surfaces^93^ and central moments were averaged within each parcels, excluding outlier vertices with values more than three scaled median absolute deviations away from the parcel median.

The BigBrain gradient was obtained from the original publication^26^ and mapped to the Desikan-Killiany^1^, Glasser^88^, and Schaefer^89^ parcellations. In brief, the authors derived an MPC matrix by correlating BigBrain intensity profiles between every pair of regions in a 1,012 cortical node parcellation, controlling for the average whole-cortex intensity profile. The MPC matrix was thresholded row-wise to retain the top 10% of correlations and converted into a normalized angle matrix. Diffusion map embedding^94, 95^, a nonlinear manifold learning technique, identified the principal axis of variation across cortical areas, *i.e.*, the BigBrain gradient. In this space, cortical nodes that are strongly similar are closer together, whereas nodes with little to no intercovariance are farther apart. To allow contextualization of surface-based findings, we mapped the BigBrain gradient to the Desikan-Killiany atlas and partitioned it into five equally sized discrete bins.

### Cytoarchitectonics data and contextualization

By adapting a previously published approach^29^, we mapped the cytoarchitectonic atlas of von Economo and Koskinas^2^ to cortical surface templates. Cytoarchitectonic class labels from the original five different structural types of cerebral cortex (agranular, frontal, parietal, polar, granular) were manually assigned to each parcellation region^29^ and subsequently mapped to vertex-wise space.

To stratify cortex-wide effects according to the five cytoarchitectonic classes, the `economo_koskinas_atlas()` function maps parcellated data (*e.g.*, disease-related atrophy map on the Desikan-Killiany atlas) to vertex-wise space and iteratively averages values from all vertices within each class.

### Connectivity data for macroscale connectome models

As in prior work^20^, we selected a group of unrelated healthy adults (*n*=207; 83 males, mean age±SD=28.73±3.73 years, range=22-36 years) from the HCP dataset^19^. HCP data were acquired on a Siemens Skyra 3T and included: (*i*) T1-weighted images (magnetization-prepared rapid gradient echo [MPRAGE] sequence, repetition time [TR]=2,400ms, echo time [TE]=2.14ms, field of view [FOV]=224×224 mm^2^, voxel size=0.7×0.7×0.7 mm, 256 slices), (*ii*) resting-state functional MRI (gradient-echo echo-planar imaging [EPI] sequence, TR=720 ms, TE=33.1 ms, FOV=208×180 mm^2^, voxel size=2 mm^3^, 72 slices), and (*iii*) diffusion MRI (spin-echo EPI sequence, TR=5,520 ms, TE=89.5ms, FOV=210×180, voxel size=1.25mm^3^, *b*-value=1,000/2,000/3,000 s/mm^2^, 270 diffusion directions, 18 b0 images). HCP data underwent the initiative’s minimal preprocessing^96^. In brief, resting-state functional MRI data underwent distortion and head motion corrections, magnetic field bias correction, skull removal, intensity normalization, and were mapped to MNI152 space. Noise components attributed to head movement, white matter, cardiac pulsation, arterial, and large vein related contributions were automatically removed using FIX^97^. Preprocessed time series were mapped to standard gray ordinate space using a cortical ribbon-constrained volume-to-surface mapping algorithm and subsequently concatenated to form a single time series. Diffusion MRI data underwent b0 intensity normalization and correction for susceptibility distortion, eddy currents, and head motion. High-resolution functional and structural data were parcellated according to the Desikan-Killiany^1^, Glasser^88^, as well as Schaefer 100, 200, 300, and 400^89^ parcellations.

Normative functional connectivity matrices were generated by computing pairwise correlations between the time series of all cortical regions and subcortical (nucleus accumbens, amygdala, caudate, hippocampus, pallidum, putamen, thalamus) regions; negative connections were set to zero. Subject-specific connectivity matrices were then *z*-transformed and aggregated across participants to construct a group-average functional connectome. Available cortico-cortical, subcortico-cortical, and subcortico-subcortical matrices are unthresholded. Normative structural connectivity matrices were generated from preprocessed diffusion MRI data using MRtrix3^98^. Anatomical constrained tractography was performed using different tissue types derived from the T1-weighted image, including cortical and subcortical gray matter, white matter, and cerebrospinal fluid^99^. Multi-shell and multi-tissue response functions were estimated^100^ and constrained spherical deconvolution and intensity normalization were performed^101^. The initial tractogram was generated with 40 million streamlines, with a maximum tract length of 250 and a fractional anisotropy cutoff of 0.06. Spherical-deconvolution informed filtering of tractograms (SIFT2) was applied to reconstruct whole-brain streamlines weighted by the cross-section multipliers^102^. Reconstructed streamlines were mapped onto the 68 cortical and 14 subcortical (including hippocampus) regions to produce subject-specific structural connectivity matrices. The group-average normative structural connectome was defined using a distance-dependent thresholding procedure, which preserved the edge length distribution in individual patients^103^, and was log transformed to reduce connectivity strength variance. As such, structural connectivity was defined by the number of streamlines between two regions (*i.e.*, fiber density).

## Acknowledgements

Many scientists contributed to the development of ENIGMA but did not take part in the writing of this report. A full list of contributors to ENIGMA is available here: http://enigma.ini.usc.edu/about-2/consortium/members/.

The authors would like to express their gratitude to the open science initiatives that made this work possible: (*i*) The ENIGMA Consortium (core funding for ENIGMA was provided by the NIH Big Data to Knowledge (BD2K) program under consortium grant U54 EB020403 to P.M.T.), (*ii*) The Allen Human Brain Atlas, (*iii*) BigBrain/HIBALL, and (*iv*) The Human Connectome Project (Principal Investigators: David Van Essen and Kamil Ugurbil; 1U54MH091657) funded by the 16 NIH Institutes and Centers that support the NIH Blueprint for Neuroscience Research; and by the McDonnell Center for Systems Neuroscience at Washington University.

## Author contributions

*Core developers*: S.L., B.C.B.

*Toolbox beta testing*: B.-Y.P., J.R., C.P., S.L.V., Y.W, M.K.

*Writing*: S.L., B.C.B.; revised and approved by other listed co-authors.

## Supplementary Information

**SUPPLEMENTARY TABLE 1.**
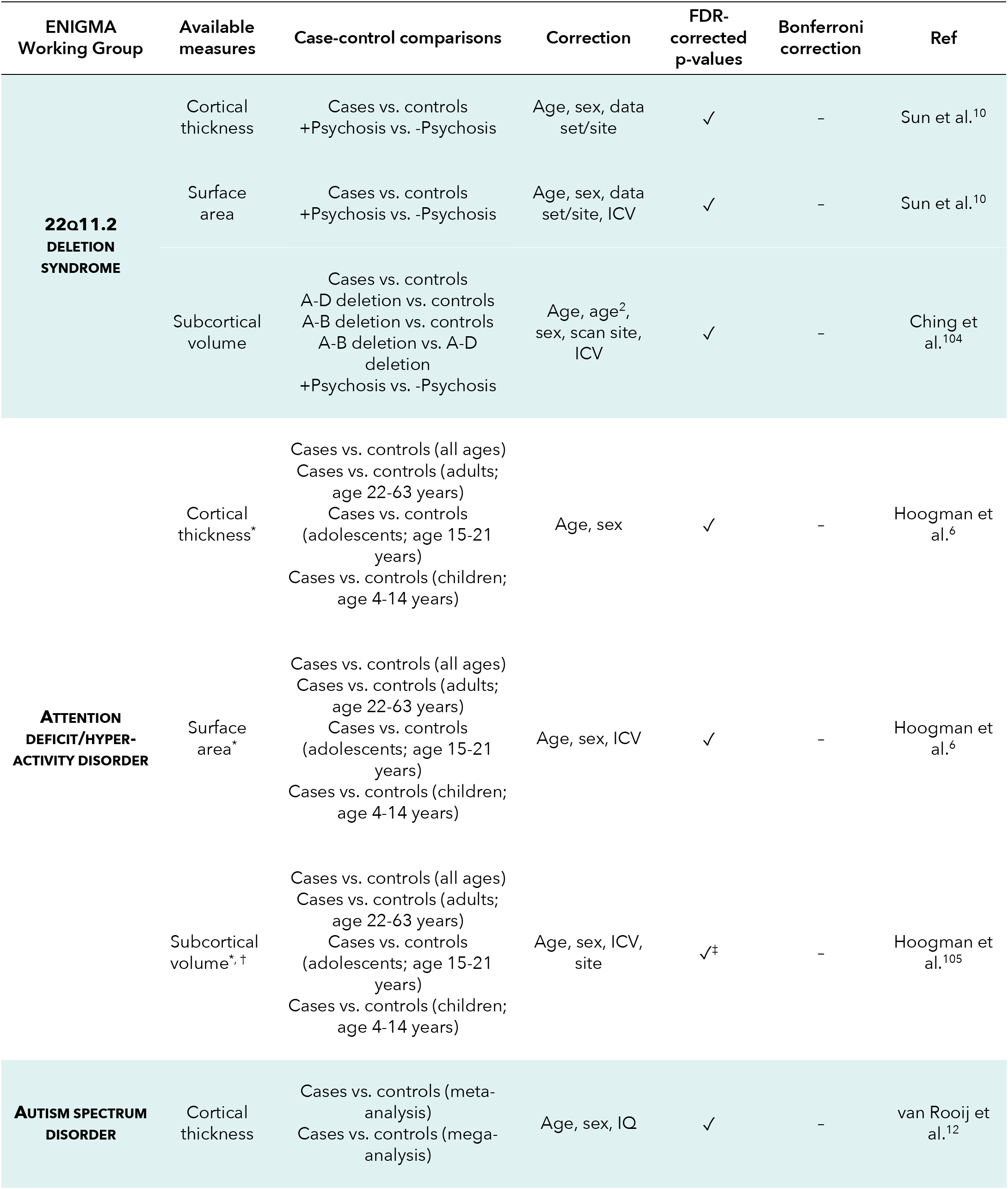
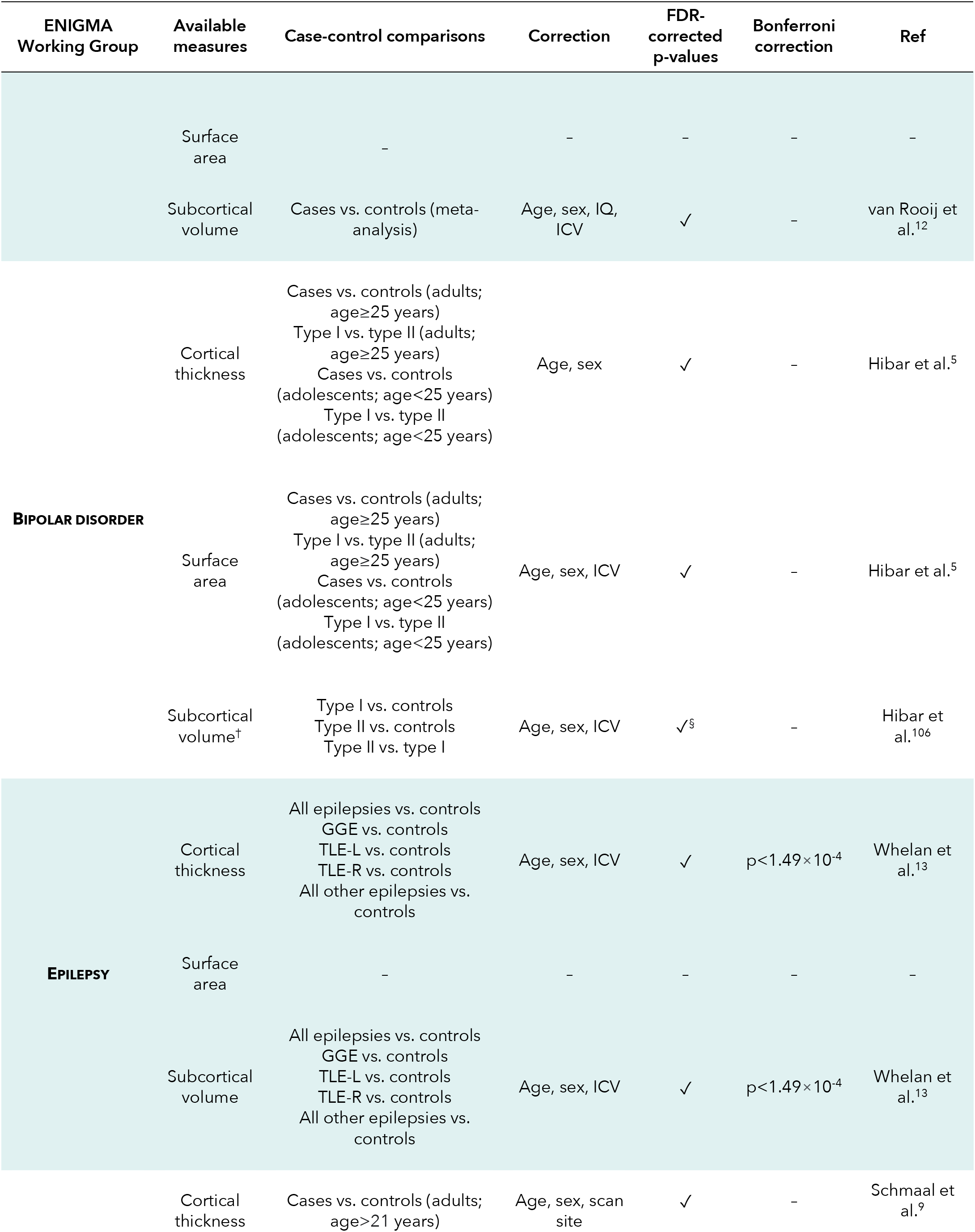
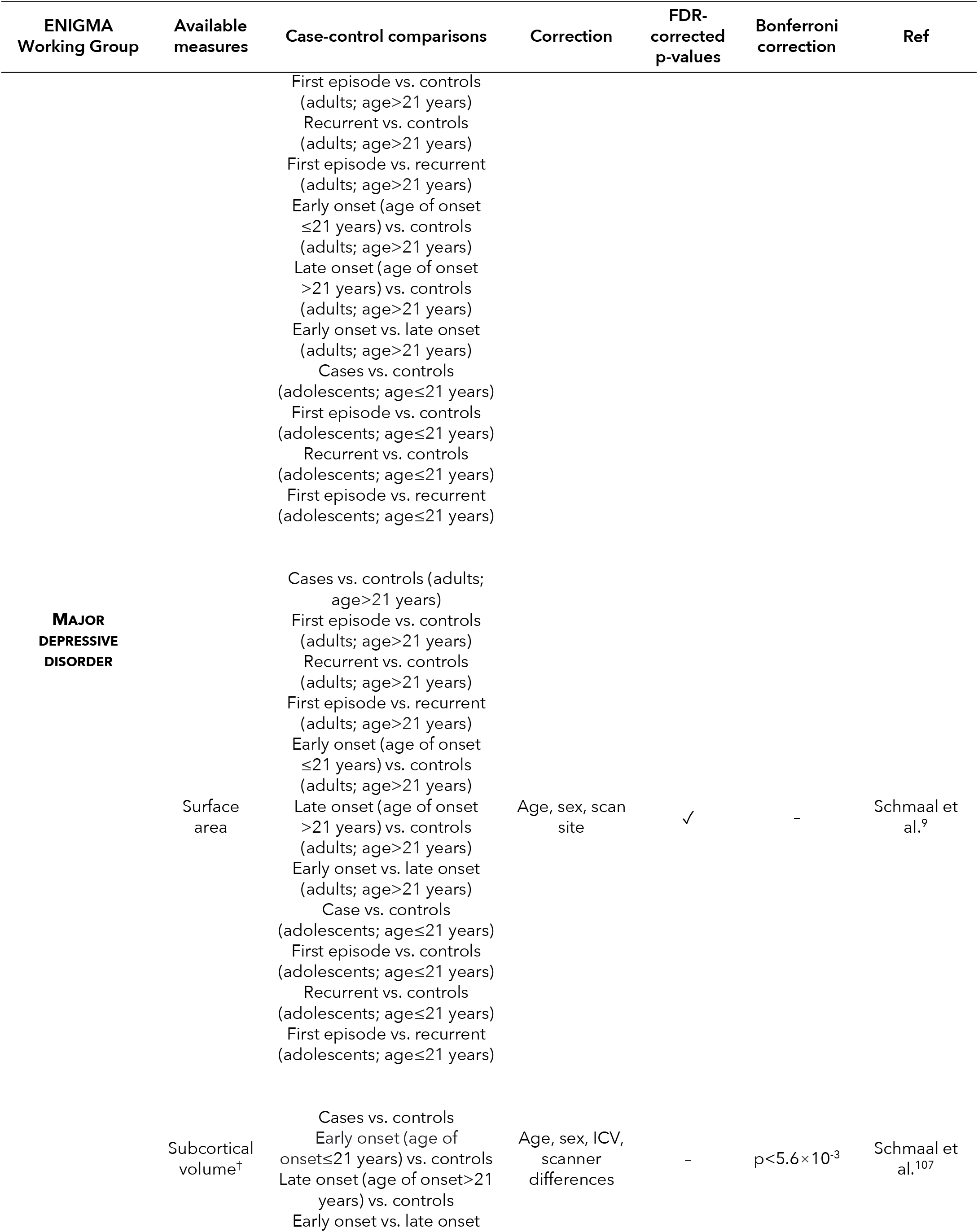
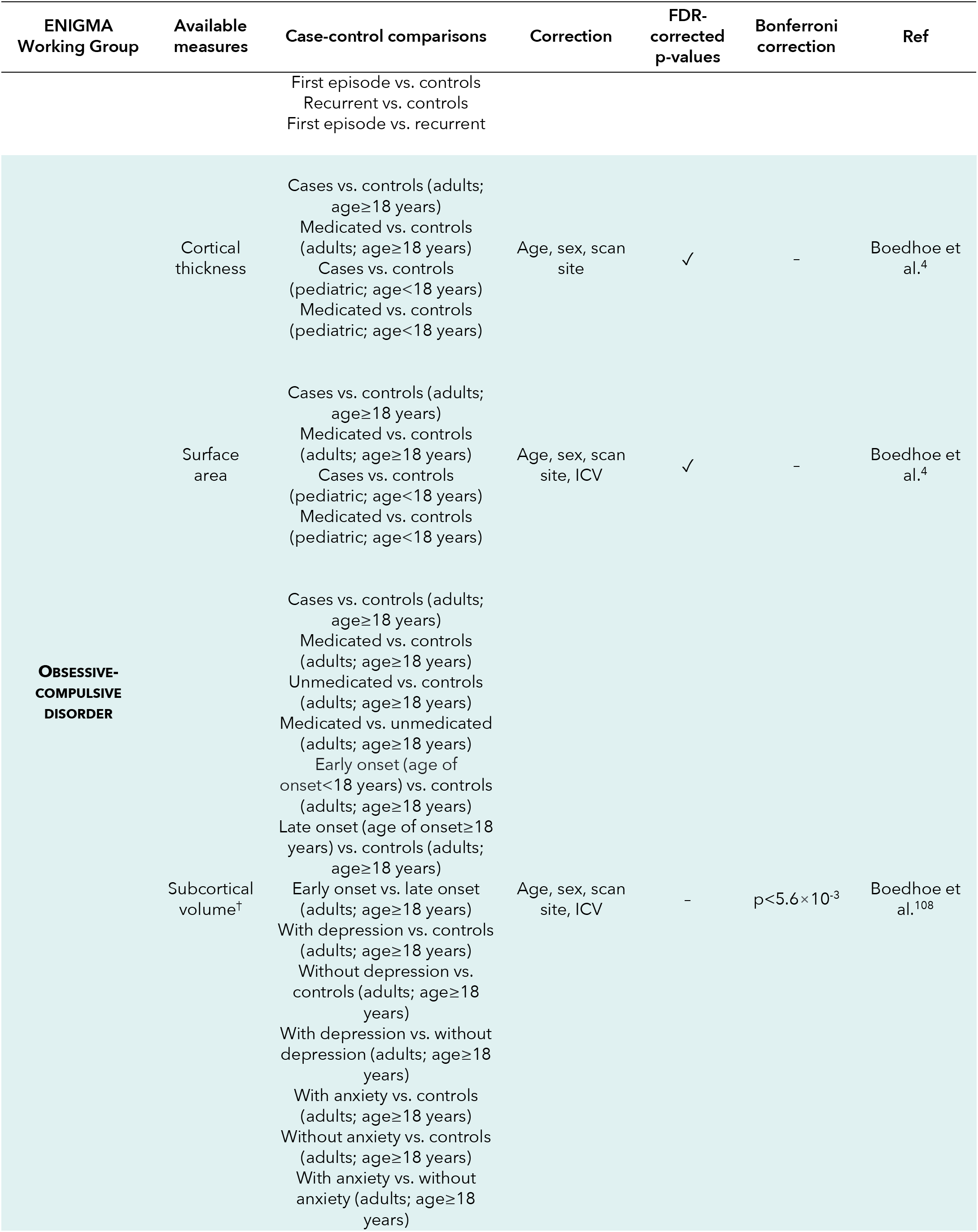
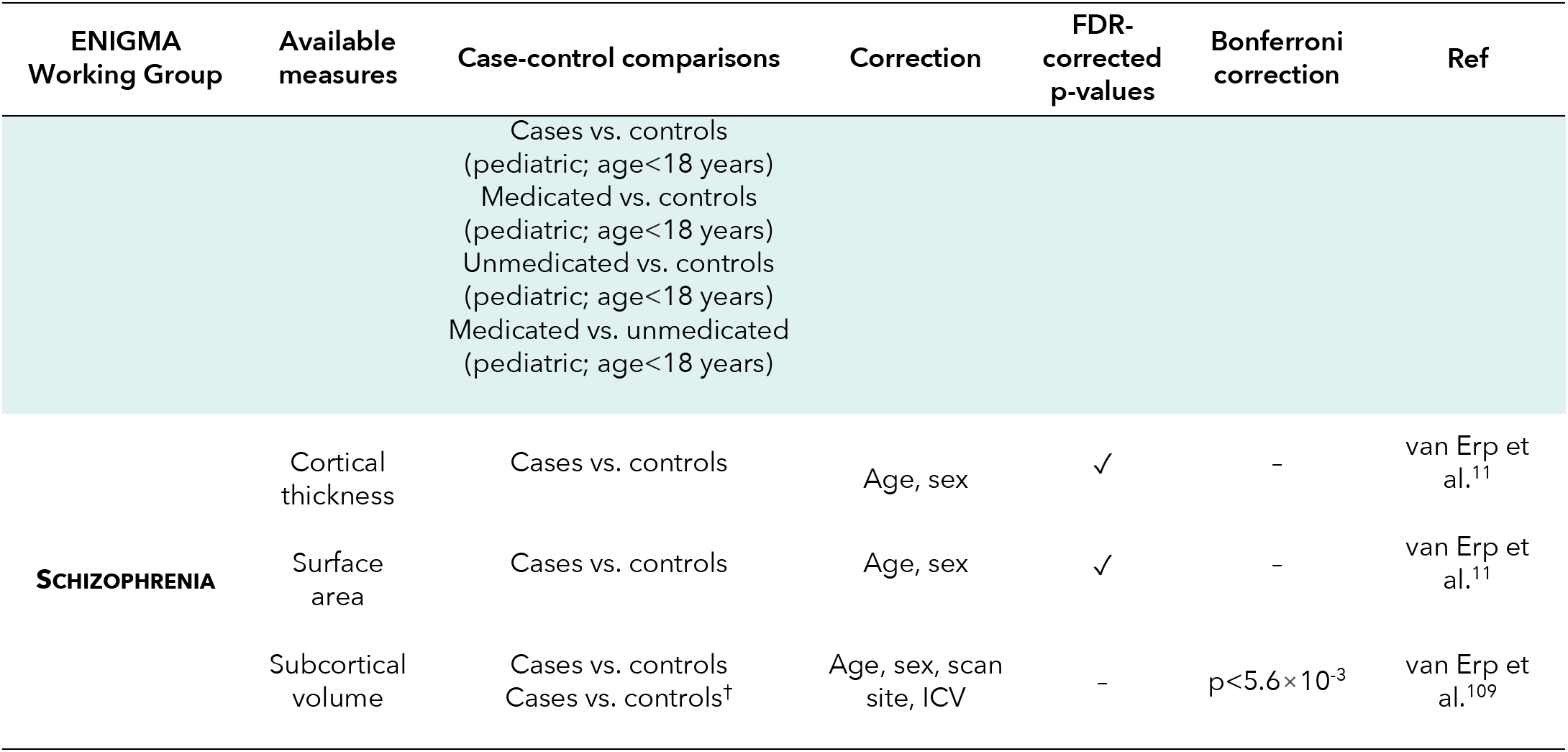
ENIGMA data description. Available case-control summary statistics. Abbreviations: FDR, false discovery rate; ICV, intracranial volume; IQ, intellectual quotient; GGE, idiopathic/genetic generalized epilepsy; TLE-L, left temporal lobe epilepsy; TLE-R, right temporal lobe epilepsy. *mega-analysis, †mean [(left+right)/2] region of interest volume, ^‡^*p*<0.0156 for FDR correction at *q*=0.05, ^§^*p*<0.00491 for FDR correction at *q*=0.05.

**FIGURE S1.**
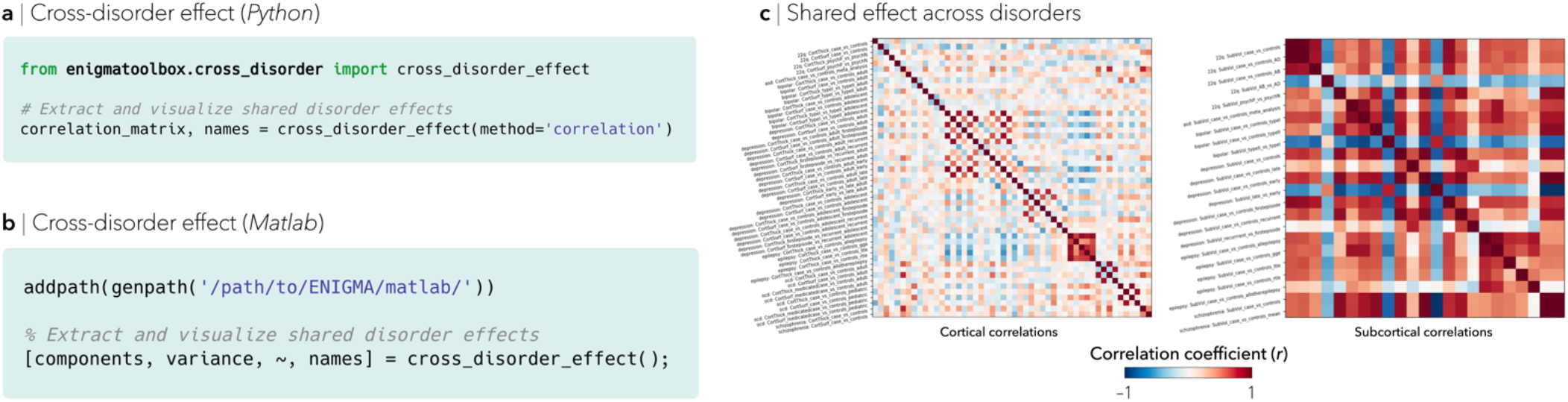
Cross-disorder effect: cross-correlation. Minimal (**a**) Python and (**b**) Matlab code snippets to systematically correlate patterns of brain structural abnormalities across disorders. (**c**) Resulting cross-correlation matrix showing similar (red) and dissimilar (blue) transdiagnostic morphometric signatures.

**FIGURE S2.**
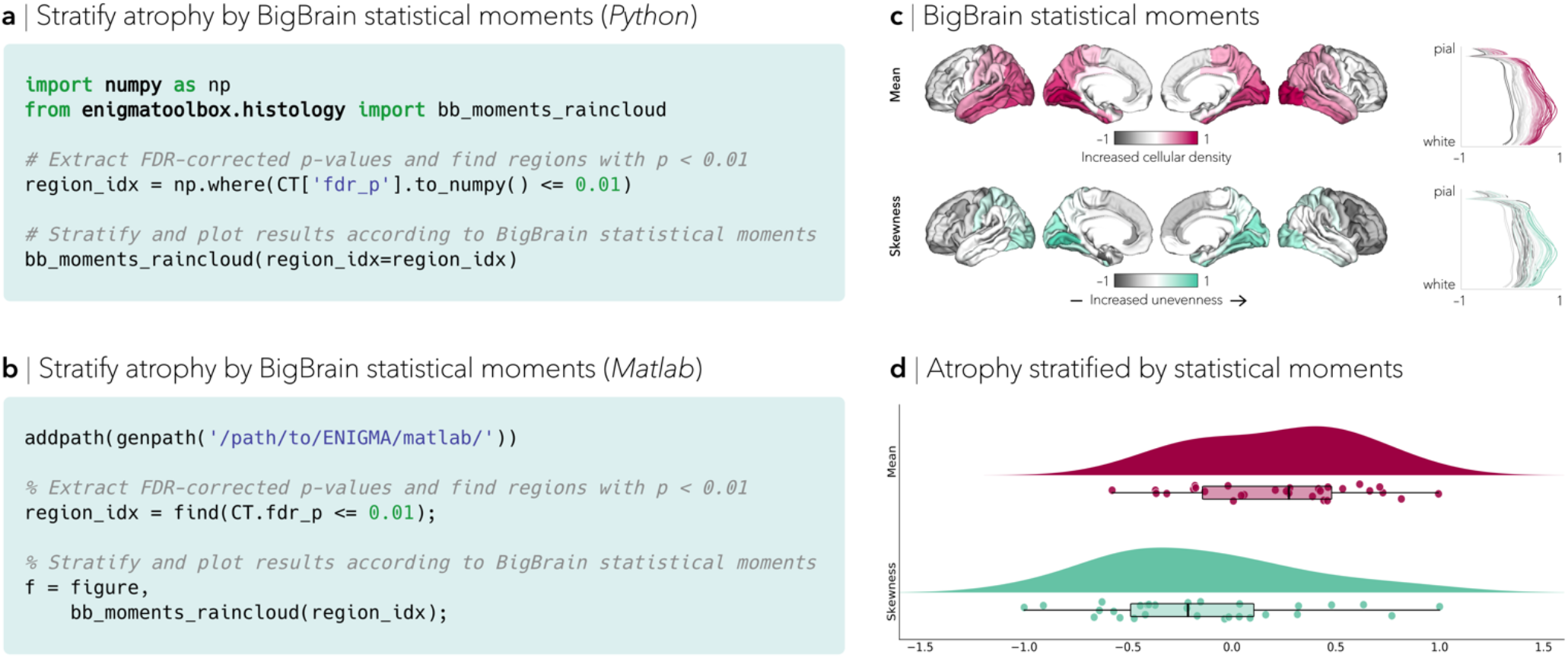
Advanced analytical workflows: BigBrain statistical moments. Minimal (**a**) Python and (**b**) Matlab code snippets to stratify significantly atrophied regions according to BigBrain statistical moments. (**c**) Regional cytoarchitecture using two statistical moments of staining profiles: (*i*) mean intracortical staining across the mantle, which allows inferences on overall cellular density, and (*ii*) profile skewness, which indexes the distribution of cells across upper and lower layers of the cortex. (**d**) Applying this approach to individuals with left focal epilepsy revealed greater cortical atrophy in regions with greater, and more evenly distributed, cellular densities.

**FIGURE S3.**
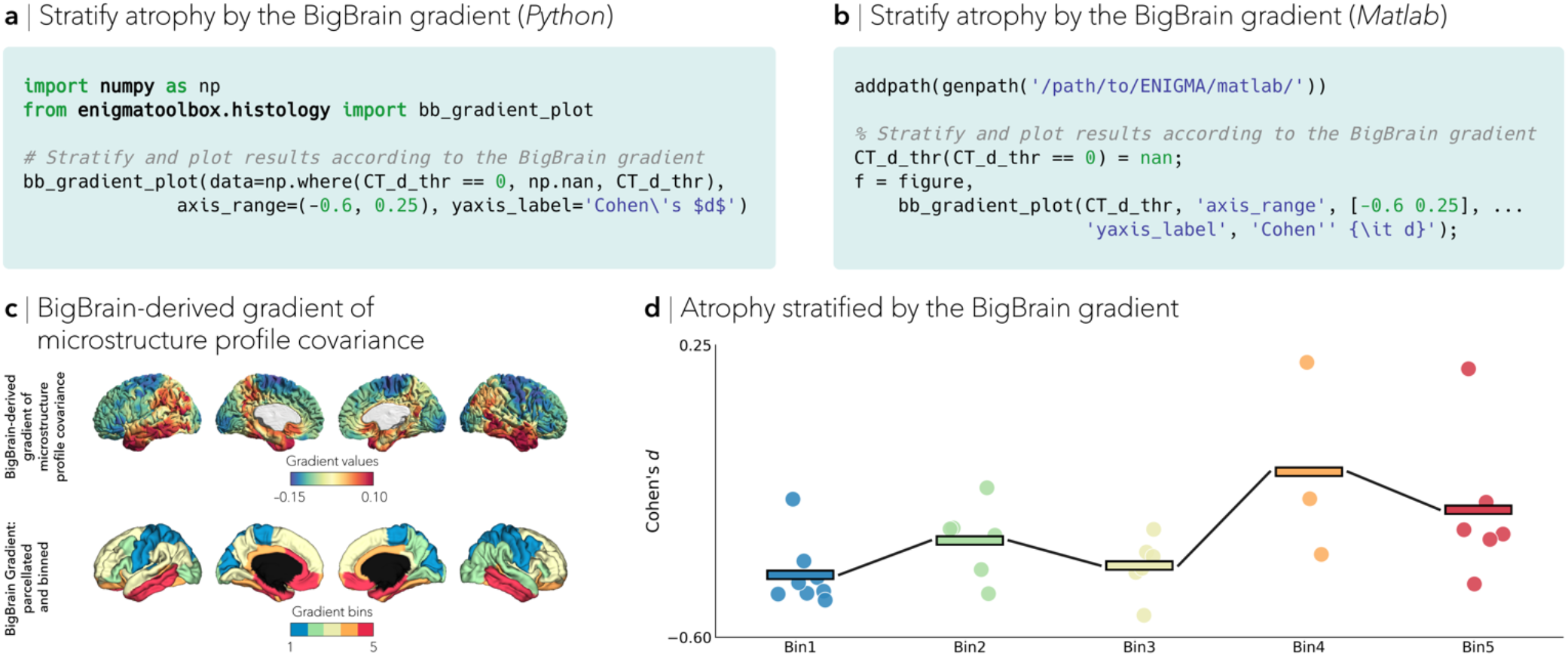
Advanced analytical workflows: BigBrain gradient. Minimal (**a**) Python and (**b**) Matlab code snippets to stratify significantly atrophied regions according to the BigBrain gradient. (**c**) BigBrain-derived gradient of microstructural profile covariance gradient describes a sensory-fugal transition in intracortical microstructure (*top*). The BigBrain gradient was then mapped to several parcellations and partitioned it into five equally sized discrete bins (*bottom*). (**d**) Applying this approach to individuals with left focal epilepsy revealed greater cortical atrophy in areas located towards the sensory apex (blue/green) of the cytoarchitectural gradient.

**FIGURE S4.**
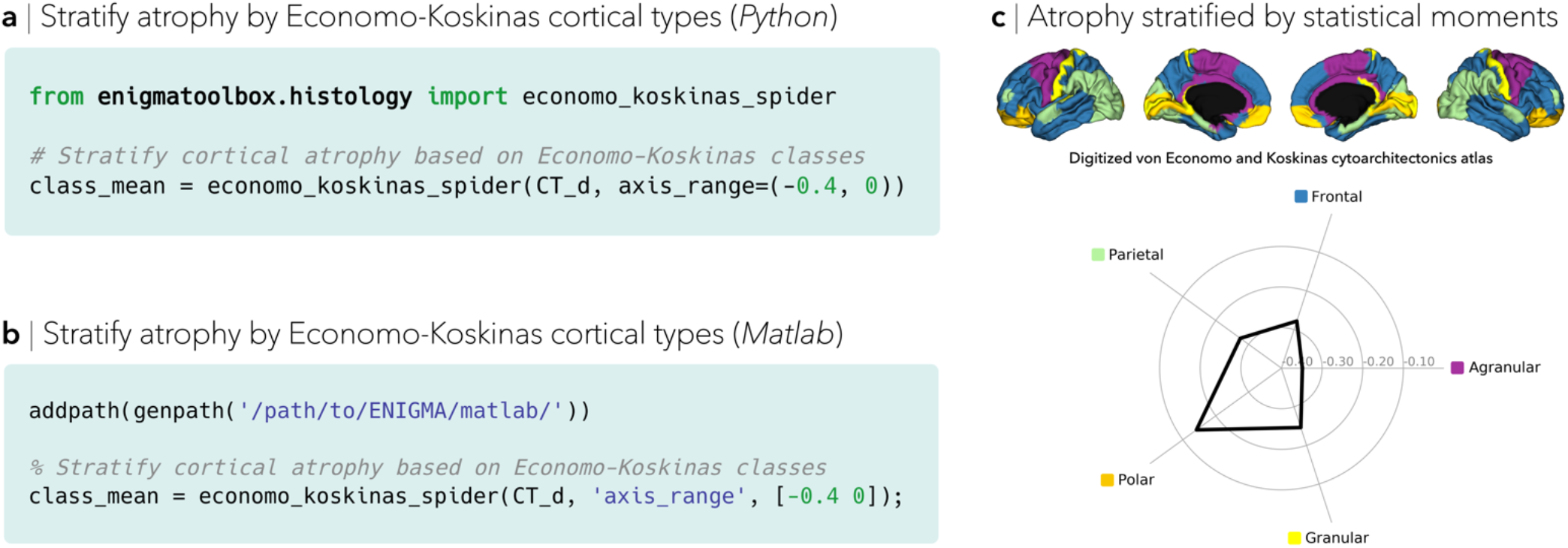
Advanced analytical workflows: von Economo and Koskinas cytoarchitectonic atlas. Minimal (**a**) Python and (**b**) Matlab code snippets to stratify significantly atrophied regions according to cytoarchitectonic classes. (c) A digitized cytoarchitectonic atlas from seminal postmortem work by von Economo and Koskinas^2^. Contextualizing cortical atrophy patterns in individuals with left focal epilepsy revealed greater atrophy in the agranular cortex (purple).

**FIGURE S5.**
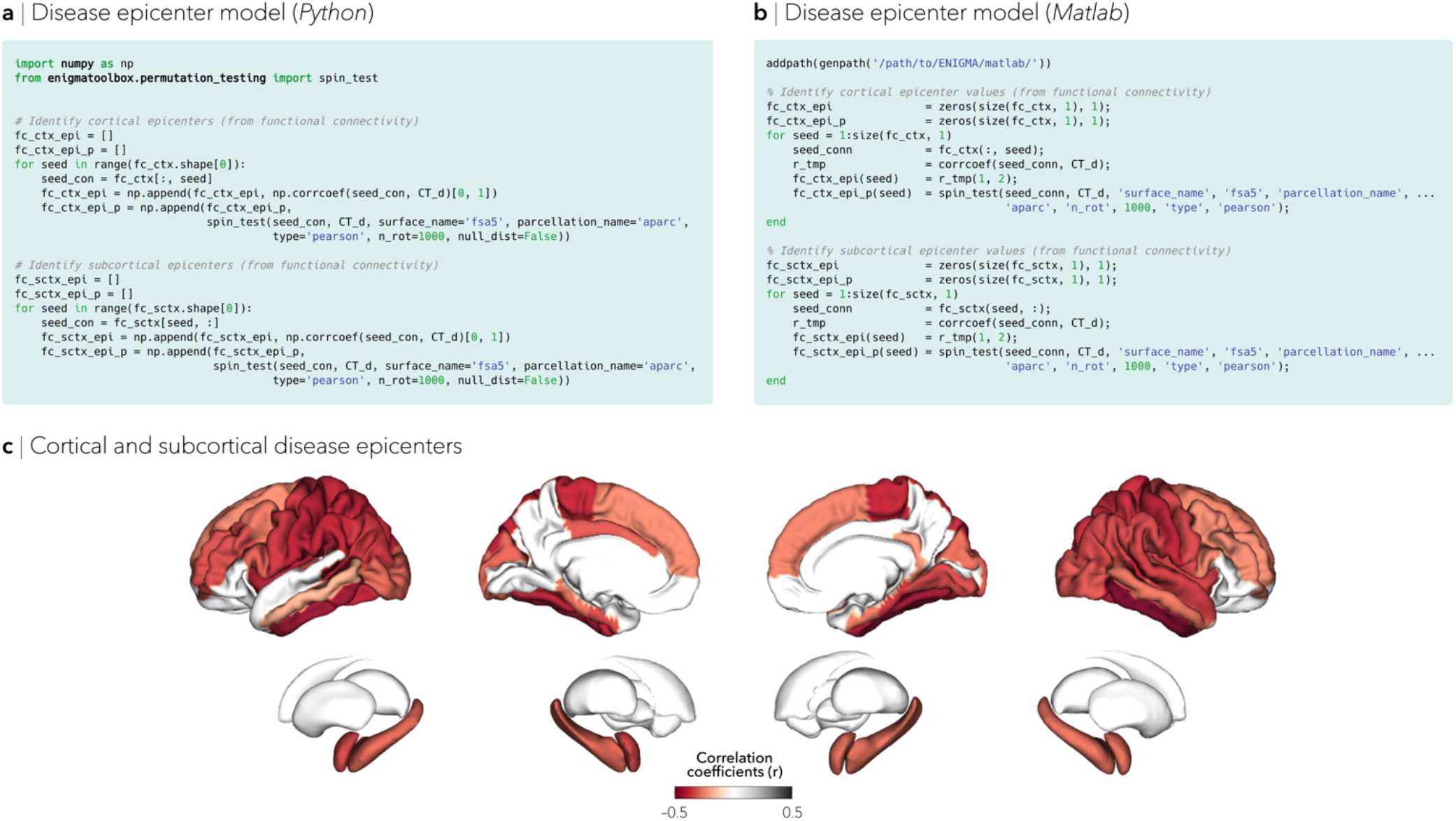
Advanced analytical workflows: disease epicenter model. Minimal (**a**) Python and (**b**) Matlab code snippets to identify disease-specific cortical and subcortical epicenters. (**c**) Applying this approach to individuals with left focal epilepsy revealed that patterns of atrophy in left focal epilepsy were anchored to the connectivity profiles of mesiotemporal regions.

